# Shedding light on cuticle formation: phytochrome B and downstream signaling events controlling cuticle deposition in tomato fruits

**DOI:** 10.1101/2025.09.11.675555

**Authors:** Leticia Danielle Longuini Gomes Fernandes, Bruna Orsi, Juliene dos Reis Moreira, Pedro Prudente do Amaral Oliveira, Jessica Naomi Motubu Ueda, Diego Demarco, Sónia Cristina da Silva Andrade, Christophe Rothan, Magdalena Rossi, Luciano Freschi

## Abstract

The cuticle is a hydrophobic barrier limiting water loss and pathogen entry. Although light influences cuticle formation, the underlying mechanisms remain poorly understood, particularly in fruits. Here, we show that fruits from tomato (*Solanum lycopersicum*) plants grown under far-red-enriched light conditions displayed increased cuticle load due to the up-regulation of cutin and wax biosynthetic genes. Both tomato PHYB-encoding genes (*SlPHYB1* and *SlPHYB2*) were implicated as negative regulators of fruit cuticle formation, as revealed by the thinner cuticles, reduced abundance of cutin monomers and cuticular waxes, and downregulation of cuticle-related genes detected in fruits from transgenic plants overexpressing constitutively active alleles of *SlPHYB1*/*B2* (*SlYHB1^OE^* and *SlYHB2^OE^*). The impaired cuticle of *SlYHB1/B2^OE^* fruits resulted in increased susceptibility to *Botrytis cinerea*, despite their markedly enhanced flavonoid accumulation in the peel. PHYTOCHROME-INTERACTING FACTORs (SlPIF3, SlPIF4) and the B-box protein SlBBX28 were identified as downstream factors of SlPHYB1/B2-mediated regulation of cuticle deposition, synergistically activating the expression of cuticle-related genes. Additionally, SlPHYB1/B2 signaling also downregulated multiple central carbon metabolism-related genes in the fruit peel, likely reducing the pool of activated fatty acids (FAs) feeding into cutin and wax biosynthetic pathways. Together, our results uncover a novel regulatory layer of light signaling in fleshy fruits, revealing that, besides controlling fruit nutritional traits, the PHYB-PIF-BBX regulatory module also shapes the fruit cuticle formation.

**Highlights:** Phytochrome-dependent light signaling regulates cuticle deposition in tomato fruits by controlling downstream transcription factors and the supply of precursors feeding into cutin/wax biosynthetic pathways, thereby affecting postharvest water loss and pathogen susceptibility.

## INTRODUCTION

The aerial surfaces of plants are covered by a lipophilic cuticle, predominantly composed of a cutin polyester layer of C16–C18 fatty acids (FAs) (Philippe et al., 2020; Kosma et al., 2024). Waxes, derived from very-long-chain fatty acids (VLCFAs) and further modified into alkanes, aldehydes, ketones, alcohols, and esters, are either embedded within the cutin layer (intracuticular) or deposited on its surface (epicuticular), and play a central role in regulating non-stomatal transpiration (Seufert et al., 2022; Staiger et al., 2019; Jolliffe et al., 2023). In addition to reducing water loss, the cuticle acts as the first line of defense against pathogens, primarily through the physical barrier formed by the cutin (Yeats and Rose, 2013). Due to these protective roles, the cuticle plays multifaceted roles in fleshy fruits, influencing key traits such as water loss, pathogen susceptibility, firmness and cracking resistance (Isaacson et al., 2009; Lara et al., 2015). As all these traits directly impact fruit shelf life, defects in cuticle biosynthesis and deposition can significantly impact fruit quality deterioration at the postharvest stage (Lara et al., 2014).

Cuticle metabolism, particularly wax accumulation, is influenced by multiple environmental cues, including water deficit, temperature, and light (Trivedi et al., 2019). Among them, light has been shown to either promote or repress cuticle formation in vegetative tissues (Hooker et al., 2002; Ma et al., 2024). In *Arabidopsis thaliana*, cuticular wax biosynthetic genes are induced in etiolated seedlings upon exposure to light (Hooker et al., 2002) and repressed in the stems of adult plants transferred to continuous darkness (Go et al., 2014). In contrast, cutin biosynthetic genes and cutin deposition in *A. thaliana* etiolated seedlings were significantly repressed soon after transferring to continuous light (Ma et al., 2024). Cuticle deposition is also altered under high light conditions (Reed and Tukey, 1982; Baker, 1974; Herzig et al., 2025), whereas the impacts of light spectral quality on cuticle formation have been only marginally investigated (Huang et al., 2020).

An array of photoreceptors connects the surrounding light information and numerous plant developmental processes (Lau and Deng, 2010). Among them, PHYTOCHROMES (PHYs) are red (R)/far-red (FR) reversible photoreceptors, existing as cytosolic inactive Pr form that is converted to active nuclear-localized Pfr upon R light absorption (Butler et al., 1959; Chen et al., 2003). In the nucleus, PHYs influence the protein stability and activity of downstream transcription factors (TFs), including the central positive or negative regulators of photomorphogenesis ELONGATED HYPOCOTYL 5 (HY5) and PHYTOCHROME-INTERACTING FACTORs (PIFs), respectively. Other photomorphogenesis-related TFs, including multiple members of the B-box protein family (BBX) and proteins of the COP1-SPA/DET ubiquitin-proteasome complex, are also integral parts of the PHY-dependent light signal transduction (Cao et al., 2023).

PHYs have recently been described as repressors of cuticular wax accumulation in maize, as loss-of-function mutants for these photoreceptors displayed increased levels of total alkanes and total free fatty acids in developing leaves (Qiao et al., 2020). In line with these findings, constitutively active *PHY* alleles, which lead to *co*nstitutive *photomorphogenic* (*cop*) phenotype, negatively impacted the expression of cutin biosynthetic genes in *A. thaliana* seedlings (Ma et al., 2024). Moreover, dark-grown seedlings of the *cop1-4 A. thaliana* mutant showed significant downregulation of genes involved in cutin biosynthesis (Ma et al., 2024). Whether similar regulatory mechanisms regulate cuticle formation in other organs, such as fleshy fruits, and the identification of other downstream components of PHY-mediated cuticle regulation remain to be explored.

PHYs are encoded by small multigene families. In tomato, a major crop and important model system for fleshy fruits, two B-type PHY-encoding genes (*SlPHYB1* and *SlPHYB2*) play unique and redundant functions in controlling fruit chloroplast biogenesis (Bianchetti et al., 2018; Alves et al., 2020), methylation reprogramming of master ripening-associated TFs (Bianchetti et al., 2022), and several metabolic pathways required for the accumulation of nutritional compounds (Bianchetti et al., 2018; Alves et al., 2020). SlPHYBs, alongside the downstream players SlHY5, SlPIFs and SlBBXs, have been progressively identified as targets for tomato fruit biofortification due to their significant impact on the levels of antioxidants, such as carotenoids, flavonoids, anthocyanins, tocopherols and ascorbate (Alba et al., 2000; Alves et al., 2020; Bianchetti et al., 2020, 2022; Rosado et al., 2019; Gramegna et al., 2019; Cruz et al., 2018; Liu et al., 2004; Llorente et al., 2016; Shiose et al., 2024; Yan et al., 2023; Zoratti et al., 2014). For example, ripe fruits of transgenic tomato plants expressing the mutant hyperactive *SlPHYB2^Y252H^* allele under the control of a fruit-specific promoter overaccumulated a wide range of health-promoting bioactive molecules (Alves et al., 2020). Moreover, this light-independent phytochrome allele significantly reduced the penalties to fruit quality due to off-vine ripening under constant darkness (Alves et al., 2023).

In contrast, knowledge on the light-dependent regulation of cuticle formation in fleshy fruits is currently limited to a few reports on postharvest treatments with monochromatic light (Giordano et al., 2023; Jaramillo et al., 2021; Cozmuta et al., 2016; Prajapati et al., 2021). Due to its thick, astomatous, and easy-to-peel cuticle, tomato fruits have been extensively used as a model for exploring the molecular mechanisms behind cuticle biosynthesis and structure, as well as their consequences for fruit shelf life, fungal resistance, and cracking (Petit et al., 2021). Using tomato fruit as a biological model, here we show that FR-enriched light conditions promote cuticle biosynthesis in fruits, suggesting a negative role for PHY-dependent light signaling in cuticle formation. Supporting this, both epicuticular waxes and cutin deposition were drastically reduced in fruits from transgenic lines overexpressing the mutant hyperactive *SlPHYB1^Y253H^* and *SlPHYB2^Y252H^* alleles (hereinafter referred to as *SlYHB2* and *SlYHB2*) due to the repression of genes involved in cutin and wax biosynthesis and deposition, and resulting in increased susceptibility to the opportunistic fungal pathogen *Botrytis cinerea*. Alongside other downstream events, specific members of *SlPIF* and *SlBBX* gene families were identified as mediators of SlPHYB-dependent light perception in controlling fruit cuticle metabolism in tomato fruits. Collectively, our results provide a novel mechanism by which the SlPHYBs and downstream TFs fine-tune the cuticle biosynthesis in fruits in response to changes in the environmental R/FR ratios, with significant consequences in fruit quality deterioration at the postharvest stage.

## MATERIAL AND METHODS

### Plant material

Wildtype (WT) tomato (cv. Moneymaker, MM) and phytochrome loss-of-function mutants (*Slphyb1*, *Slphyb2* and *Slphyb1b2*) were obtained from Rameshwar Sharma, University of Hyderabad, India (Appenroth et al., 2006). Transgenic tomato (cv. MicroTom, MT) lines silenced for *SlPIF4* (Solyc07g043580) and *SlBBX28* locus (Solyc12g005660) were previously obtained and characterized by Rosado et al (2019) and Lira et al (2022), respectively (named hereafter *SlPIF4^RNAi^ and SlBBX28^RNAi^*). Silencing of *SlPIF3* (*SlPIF3^RNAi^*) was obtained by constitutively expressing an intron-spliced hairpin RNA construct containing a 265-bp fragment of the *SlPIF3* locus (Solyc01g102300).

Transgenic MT lines constitutively overexpressing the Tyr^253^-to-His *SlPHYB1* mutant version (*SlYHB1^OE^*) and the Tyr^252^-to-His *SlPHYB2* mutant version (*SlYHB2^OE^*) were obtained by amplifying the full-length cDNAs encoding *SlPHYB1* (Solyc01g059870) and *SlPHYB2* (Solyc05g053410). Each fragment was inserted into entry vectors pDON221 (Thermo Scientific) and recombined with pK7WG2D (Karimi et al., 2002) using LR Clonase (Invitrogen). The single T-to-C base change required to modify the aminoacid translation from Tyr^253^ to His^253^ and Tyr^252^ to His^252^ in *SlPHYB1* and *SlPHYB2*, respectively, was inserted using the QuikChange Lightning Site-Directed Mutagenesis Kit (Agilent), resulting in the *35S::SlPHYB1^Y253H^* (named hereafter *SlYHB1^OE^*) and *35S::SlPHYB2^Y252H^* (named hereafter *SlYHB2^OE^*) constructs, respectively.

In all cases, the *Agrobacterium*-mediated transformation of tomato plants cv. Micro-Tom harboring the wildtype *SlGOLDEN2-LIKE 2* (*SlGLK2*) allele (Carvalho et al., 2011) was conducted as described by Bianchetti et al. (2018). The homozygous Micro-Tom (MT) lines were obtained from the T_2_ or T_3_ progeny by seeking 100% kanamycin resistance in the T_3_ or T_4_ seed population, respectively. Experiments were performed in homozygous T_4_ (or later) generations.

### Growth conditions

Unless otherwise specified, the tomato plants were grown under standard greenhouse conditions, as described in Cruz et al. (2018). In brief, MT and MM plants were grown in 1-L and 30-L rectangular pots, respectively, containing a 1:1 mixture of commercial substrate (Plantmax HT, Eucatex, São Paulo, Brazil) and expanded vermiculite, supplemented with 1 g L^−1^ of NPK 10:10:10, 4 g L^−1^ of dolomite limestone (MgCO_3_ + CaCO_3_) and 2 g L^−1^ thermophosphate (Yoorin Master®, Yoorin Fertilizantes, Brazil) in greenhouse under automatic irrigation at an average mean temperature of 25◦C, natural light 11 h/13 h (winter/summer) photoperiod and approximately 250–350 µmol m^−2^ s^−1^ PAR irradiance.

For experiments under controlled conditions with distinct red to far-red (R/FR) ratios, wild type plants were growth under LED lamps adjusted for a R/FR ratio of 2.3 (control condition) and low R/FR (0.23, +FR condition), always maintaining 250 µmol m^-2^ s^-1^, 12-h photoperiod, air temperature of 25 °C/18 °C day/night and 60 %/80 % day/night relative air humidity.

### Postharvest water loss measurements

For water loss measurement, approximately 30 fruits of each genotype were harvested at the ripe (Breaker plus 7 days, Bk7) stage and kept at room temperature for up to 2 months. Fruit weight was recorded periodically, and water loss was calculated as a percentage of the weight loss.

### Epicuticular wax analysis

Gas chromatography–flame ionization detector (GC-FID) and GC coupled to a mass spectrometer (GC-MS) combined profiling of epicuticular waxes was conducted as described by Fernandez-Moreno et al (2016), with some modifications. The epicuticular waxes were extracted by dipping intact ripe fruits in chloroform for 1 min, spiked with 100 µg tetracosane as an internal standard. The samples were then completely evaporated, resuspended in 50 µL pyridine and derivatized with 50 µL bis-*N,O*-(trimethylsilyl) trifluoroacetamide (BSTFA) at 80 °C for 60 min under agitation (1200 rpm). Sample volumes of 1 mL of the trimethylsilylated extracts were analyzed by GC-FID (Trace GC Ultra gas chromatography, Thermo Electron). The chromatograph was equipped with a fused-silica capillary column (30 m, ID 0.25 mm, 0.25 mm thick internal film) DB-5 MS stationary phase using nitrogen as the carrier gas at a flow rate of 1.25 mL min^−1^. Injector and detector temperatures were 300 and 250 ^°^C, respectively. Separation of cuticular waxes was performed using a temperature program of 1 min at 150 °C, raised by 10 °C min^−1^ to 250 °C, subsequently increased by 3 °C min^−1^ to 320 °C, and held for 7 min at 320 °C.

Wax constituents were identified by GC-MS (model QP2010 SE, Shimadzu), with total ion current spectra recorded in the mass range of 50–700 atomic mass units in scanning mode. The injector temperature was 300 ^°^C, and the following mass spectrometry (MS) operating parameters were used: ionization voltage, 70 eV (electron impact ionization); ion source temperature, 230 ^°^C; interface temperature, 320 ^°^C. Single compounds were quantified against the internal standard by manually integrating peak areas. Compounds with a proportion < 0.01 % of the total wax were included in the data analysis. Data were expressed as ug cm^-2^, and the total surface area of tomato fruits was calculated from the average of vertical and horizontal diameters by assuming a spherical shape.

### Cuticle isolation and cutin monomer analysis

Cuticles were enzymatically isolated from ripe tomato fruits according to (España et al., 2014b), using a solution of citrate buffer (50 mM; pH 3.7) containing 0.2 % (w/v) fungal cellulase, 2.0 % (w/v), and 1 mM NaN_3_ to prevent microbial growth. Samples were incubated at 30 °C for at least 24 h, and the cuticle was then separated from the epidermis, rinsed in distilled water, dried at 50 °C for 3 d, and stored under dry conditions. The cuticle surface area was calculated using ImageJ software, and the samples were then weighed for use in the cutin monomer analysis.

The cutin monomer profiling of enzymatically-isolated cuticular membranes was performed following a protocol based on Jenkin and Molina (2015), with some modifications. Briefly, the isolated cuticular membranes were delipidated in a methanol: chloroform (2:1 v/v) mixture overnight at room temperature under agitation (100 rpm). Approximately 10 mg of dewaxed cuticular membranes were transferred to a tube and added with 100 mg methyl heptadecanoate as an internal standard. A 6-mL reaction mixture, consisting of 0.9 mL methyl acetate, 1.5 mL sodium methoxide and 3.6 mL methanol, was added to each sample. The vials were capped and heated to 60 °C for 4 h. After cooling down to room temperature, 10 mL diethyl ether, 1.5 mL glacial acetic acid and a saline solution (0.5 M NaCl) were added to fill each tube, and the tubes were mixed thoroughly by vortexing for 1 min. Samples were centrifuged for 10 min at 1000 *g,* and the upper organic phase was removed to another tube. Saline solution (0.5 M NaCl) was added to fill each tube, vortexed for 1 min, and centrifuged for 10 min at 1000 *g*. The aqueous upper phase was removed, and the washing procedure with saline solution was repeated. After the removal of aqueous upper phase, anhydrous sodium sulfate was added to the solvent, and the tubes were vortexed for 1 min and then incubated overnight. Samples were centrifuged for 2 min at 1000 *g,* and the organic phase was transferred to a separate tube. The solvent was evaporated before derivatization with 50 µL pyridine and 50 µL BSTFA at 80 °C for 60 min.

GC-FID and GC-MS combined profiling of cutin monomers was perfomed by anayzing 1 µL sample volumes of the trimethylsilylated extracts via both GC-FID and GC-MS as used for the separation and identification of cuticular waxes running the temperature program of 1 min at 100 °C, raised by 15 °C min^−1^ to 230°C, then increased by 5 °C min^−1^ to 240 °C, subsequently increased by 20 °C min^−1^ to 320 °C and held for 6 min at 320 °C. Cutin monomers were identified in the total ion chromatogram by comparing their mass spectra with those of authentic standards and data from the literature.

### Flavonoid Profiling

Flavonoids were profiled in the isolated peel and flesh tissues of ripe fruits as described in Alves et al. (2020), with some modifications. Approximately 20 and 250 mg fresh weight (FW) of powdered peel and flesh samples, respectively, were extracted with 0.5 mL of 80 % (v/v) methanol containing 50 µg morin as an internal standard. After incubation for 30 min in an ultrasonic bath at room temperature, the supernatant was collected by centrifugation (13 000 *g*, 2 min, 25 °C). Compounds were identified by HPLC-DAD 1260 system (Agilent Technologies, USA) equipped with an autosampler, using a Zorbax Eclipse Plus C18 column (100 x 4.6 mm, 3.5 µm particle diameter) at 45 °C with a flow rate of 1 mL min^-1^. The mobile phase was a gradient of 0.1 % acetic acid (A) and acetonitrile (B): 0 to 10 min with 100 % to 85 % A, 10 to 24 min 85% to 70%, 24 to 30 min 70% to 0% A. Eluted compounds were detected and quantified at 280 and 352 nm wavelengths. The endogenous metabolite concentration was obtained by comparing the peak areas of the chromatograms with commercial standards.

### Microscopy

For the light microscopic imaging of outer pericarp sections, ripe fruits were sliced into sections of 0.5 cm^2^ and fixed in a FAA (formalin–acetic acid–ethanol 50 %, 1:1:18) for 24 h under vacuum. Samples were then dehydrated in a graded butanol and embedded in Paraplast Plus (Sigma). Sections (12 µm thick) were obtained by using a rotary microtome and stained with Sudan IV. The distance from the cuticle outer surface to the edge of the epidermal cell (cuticle thickness), the depth of cell wall cutinization between epidermal cells (anticlinal peg depth), and the distance from the outer surface to the point of its deepest penetration into the tissue (invagination degree) were determined using a light microscope (Leica® DM 4000 B).

### *Botrytis cinerea* inoculation assay

*Botrytis cinerea* (strain B05) was grown and maintained on potato dextrose agar medium (1.5% agar, 2% potato extract, 2% dextrose). Conidia were harvested from agar plates using approximately 5 mL of the germination solution (189 mM dextrose and 70 mM KH_2_PO_4_) and a glass rod, filtered, and then resuspended in approximately 5 mL of the germination solution (Elad, 1991). Briefly, on the day of harvesting, ripe fruits were disinfected with 10% (v/v) commercial bleach, followed by three rinses with sterile water, and then inoculated by applying 5 µL of the spore suspension (1,2 x 10^6^ spores mL^-1^) onto the fruit surface at four equidistant points around the equatorial region. As a negative control, a mock inoculation with 5 µL of the suspension medium (189 mM dextrose and 70 mM KH_2_PO_4_) was applied to a single site at the base of the fruit. After inoculation, the fruits were incubated for 3 d at 25 °C in a dark and damp environment. Susceptibility was monitored by assessing disease incidence (percentage of affected lesions) and severity (lesion area). The lesion area was quantified by photographing the lesions and processing the images using ImageJ software (Schneider et al., 2012).

### Yeast two-hybrid assay

To confirm protein-protein interaction between SlBBX28 and SlPIF3 or SlPIF4, we performed the yeast two-hybrid (Y2H) assay. The full-length open reading frames with stop codon (ORFs) of *SlBBX28*, *SlPIF3* and *SlPIF4* were PCR-amplified from leaf cDNA using specific primers (Supplementary Table S1) and introduced into the pCR™ 8/GW/TOPO TA Cloning (Invitrogen) vector. Further, the ORFs were recombined into both pGADT7 or pGBT9 Gateway™ destination vectors, at the N-terminus of the activation domain (AD) or DNA-binding domain (BD) of the *GALACTOSIDASE 4* (*GAL4*) transcriptional activator, respectively (Cuéllar et al., 2013), using LR Clonase II enzyme mix (Invitrogen). *Saccharomyces cerevisiae* PJ69-4A lines (James et al., 1996) were co-transformed with both destination vectors using the polyethylene glycol (PEG)/lithium acetate method. Transformants were selected on SD (synthetic minimal) medium (Takara Bio, Shigo, Japan) lacking leucine and tryptophan. Three individual colonies were grown overnight in liquid cultures at 30 °C, and 10-or 100-fold dilutions were dropped on control (SD-Leu-Trp) and selective media (SD-Leu-Trp-His).

### Gene promoter analysis

Gene promoter analysis was performed using the promoter sequences available at the Sol Genomics Network. Typically, 3 Kb upstream of the initial ATG codon of each sequence was analysed using the PlantPAN 2.0 platform (http://plantpan2.itps.ncku.edu.tw/) (Chow et al., 2016) for the presence of PBE-box (CACATG), G-box (CACGTG - TACGTG), E-box (CANNTG, Zhang et al., 2013), CAAT (Ben-Naim et al., 2006), CORE2 (TGTGN2-3ATG, Tiwari et al., 2010) and ACE-box (ACGT, Wang et al., 2021) motifs.

### Transient expression assay (TEA) in Bright Yellow-2 (BY-2) cells

The effectors constructs were obtained by recombing the full-length ORF of *SlPIF3*, *SlPIF4* and *SlBBX28* into p2GW7 destination vector (Vanden-Bossche et al., 2013) using LR Clonase II enzyme mix (Invitrogen). The *SlCER1.2* (985 bp), *SlCER6* (1046 bp), *SlCER26* (1601 bp), *SlKCS10* (1235 bp), *SlGDSL* (1348 bp), *SlCUS1* (1161 bp), *SlCYP86A69* (966 bp) and *SlABCG11* (1022 bp) promoter sequences were PCR-amplified from leaf gDNA and cloned into pCR™ 8/GW/TOPO TA Cloning vector (Invitrogen). Next, Gateway LR Clonase II enzyme mix (Invitrogen) was used to introduce the fragments into pGWL7 destination vector (Vanden-Bossche et al., 2013), obtaining the promoter constructs harboring *FIREFLY LUCIFERASE* (*fLUC*) reporter gene. TEAs were performed in protoplasts prepared from *Nicotiana tabacum* BY-2 cells, as previously described by Shiose et al. (2024). Briefly, protoplasts were co-transfected with promoter/effector combinations. For normalization, a construct with the *RENILLA LUCIFERASE* (*rLUC*) reporter gene under the control of the cauliflower mosaic virus (*CaMV*) *35S* promoter was used. A construct containing *pCaMV35S::GUS* was used as a negative effector control. fLUC activity was expressed as the fLUC/rLUC activity ratio relative to the negative control.

### Luciferase transactivation assays

Effector constructs were generated by introducing the full-length ORFs of the *SlPIF4* and *SlBBX28* genes into the pK7WG2D vector (Karimi et al., 2002) using LR Clonase II enzyme mix (Invitrogen). The promoters, previously cloned into the pCR™ 8/GW/TOPO TA Cloning vector (Invitrogen), were recombined into the pKGWL7 (Karimi et al., 2005) vector to obtain the reporter constructs.

For promoter transactivation, *Agrobacterium tumefaciens* (GV3101) harboring the reporter and the effector constructs was co-infiltrated into leaves from five-week-old *N. benthamiana* plants. After 72 h, the leaves were sprayed with 0.8 mM D-luciferin (Promega) and incubated in the dark for 15 min. Relative luciferase (LUC) activity was quantified using the NEWTON 7.0 CCD imaging system (Vilber). At least five different leaves were evaluated for each combination, and the *35S::NLS-GFP* vector was used as the effector negative control.

### RNA isolation, quantitative PCR (qPCR) analysis and RNA sequencing

Total RNA extractions were performed using ReliaPrepTM RNA Tissue Miniprep System (Promega) from manually dissected peel tissues pooled from five to eight immature green (IG, 15 dpa, d post anthesis) tomato fruits. cDNA synthesis and RT–qPCR analysis were performed as described in Quadrana et al. (2013). Transcript abundance was normalized against the geometric mean of two reference genes, *CAC* and *EXPRESSED*. Primer sequences used are presented in Supplementary Table S1.

RNA sequencing was performed in triplicate at Novogene (Novogene Co, USA). The mRNA library was constructed using the Illumina TruSeq RNA Library Prep Kit v2 (Illumina, USA) according to the manufacturer’s protocol. A total of 36 mRNA libraries were sequenced using the NovaSeq 6000 system (Novogene Co, USA) to generate 2 × 150 bp paired-end reads.

### Read mapping, transcript profiling, and functional enrichment analysis

Raw RNA-seq data were filtered for low-quality reads, primer sequences, and vectors using Seqyclean (v1.9.10), with a cutoff of bases having an average quality lower than 24QScore. The Univec database was used to filter contaminants out, and reads smaller than 65 bp were removed. Filtered reads were aligned to the tomato genome reference SL4.0, with gene models ITAG4.1 from the SOL Genomics Network database (https://solgenomics.net/ftp/tomato_genome/annotation/ITAG4.1_release/). Both alignment and counting were performed with STAR (v2.6.1) (Dobin et al., 2013).

Only the uniquely mapped reads were used for differential expression analysis. The edgeR package (v4.4.2) (Robinson et al., 2010) from the R/Bioconductor software was used to obtain the differentially expressed genes (DEGs). Within the package, the data from each group were normalized with TMM (Robinson and Oshlack, 2010) using the calcNormFactors function. Contigs with expression profiles of zero in at least three samples were removed to avoid artifacts caused by low-expression contigs. Differential expression analysis was performed using the negative binomial function, applying the Benjamini-Hochberg (BH) correction (Benjaminit and Hochberg, 1995) for multiple tests to control the False Discovery Rate (FDR). Only genes with FDR ≤ 0.05 were considered differentially expressed. Kyoto Encyclopedia of Genes and Genomes (KEGG) enrichment analysis was performed with the R package ClusterProfiler (v4.4.16) on a pre-ranked gene list based on |log2FC| (Xu et al., 2024). The gseKEGG function was applied with the following parameters: pvalueCutoff = 0.05, pAdjustMethod = BH, exponent = 0, and maximum and minimum size of gene set of 800 and 5, respectively.

### Statistical analysis

The experimental design was completely randomized. Statistical differences between groups were determined by ANOVA followed by Student’s t-test or Tukey’s HSD test. Statistical analyses were performed using IBM SPSS Statistics (v25.0, 2020).

## RESULTS

### Simulated proximity shade promotes cuticle formation in tomato fruits

Reduced R/FR ratios are typically observed under conditions of neighbor proximity shade or full shade, and influence multiple aspects of plant development (Fernández-Milmanda and Ballaré, 2021; Tan et al., 2022). In proximity shade, the photosynthetically active radiation (PAR, 400–700 nm) intensity remains unaffected, but FR is reflected by neighboring vegetation, whereas in full shade conditions, both R/FR ratio and PAR are reduced (Martinez-Garcia and Rodriguez-Concepcion, 2023). By comparing fruits from wildtype (WT, cv MicroTom) plants grown under simulated proximity shade, our findings reveal significant impacts of high and low R/FR ratios (R/FR of 2.3 and 0.23, respectively) on multiple cuticle-related traits (Fig. 1).

**Figure 1.**
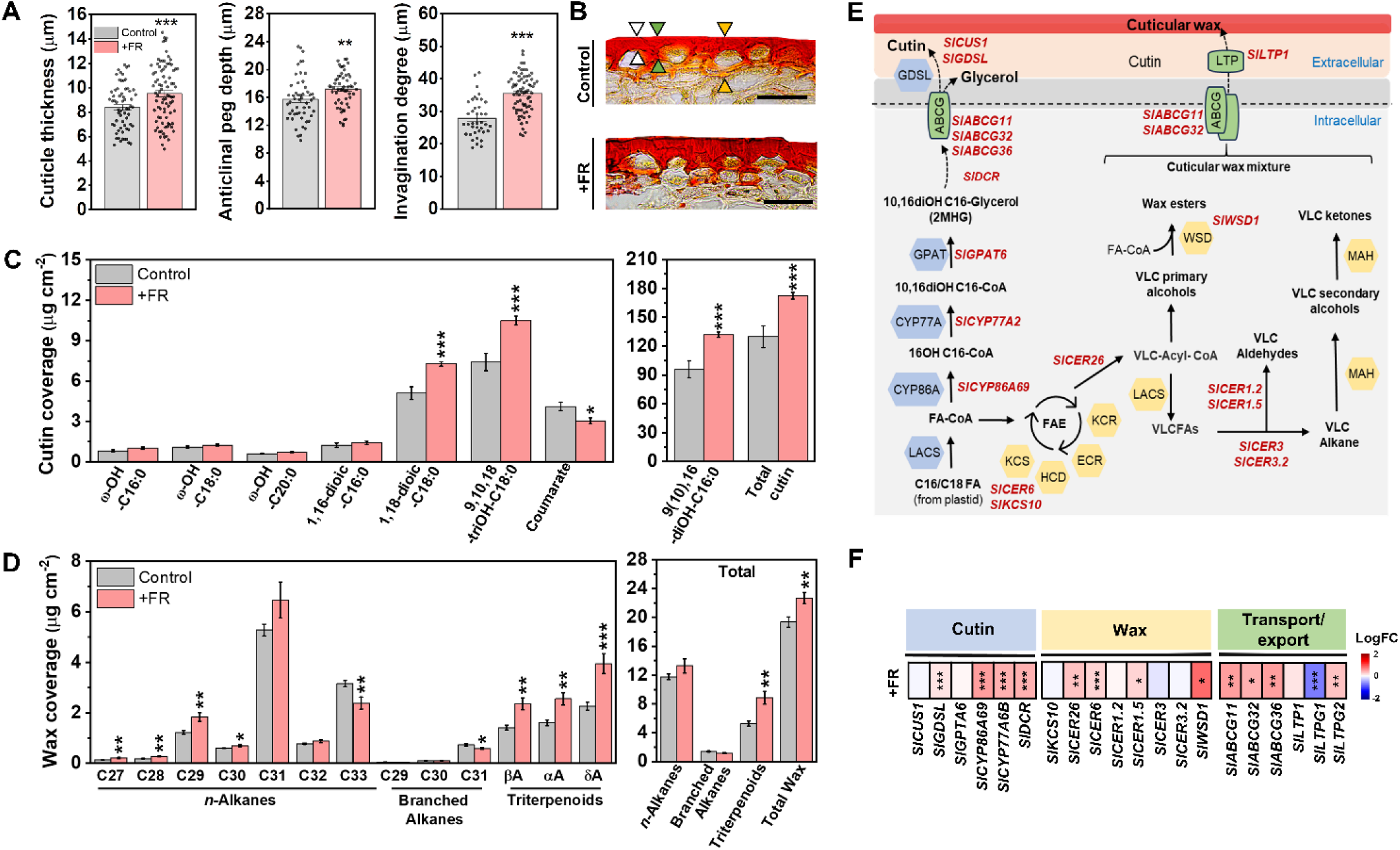
Far-red-enriched light promotes cuticle biosynthesis and deposition in tomato fruits. Measurements were performed in fruits from wildtype (WT) plants (cv. MicroTom) grown under control conditions (high R/FR, 2.3) or +FR conditions (low R/FR, 0.23). **A)** Cuticle thickness, anticlinal peg depth and invagination degree of ripe fruits (7 d after Breaker stage) (*n*=5 fruits). **B)** Representative light micrographs of pericarp cross-sections stained with Sudan-IV. White arrowheads indicate the cuticle layer over epidermal cells (reference region for cuticle thickness measurements), green arrowheads indicate the cutinization depth between epidermal cells (reference region for anticlinal peg depth measurements), and yellow arrowheads indicate the cutinized subepidermal cell walls (reference region for invagination degree). Scale bars: 50 μm. **C)** Cutin monomer composition, total cutin load of ripe fruits (*n*=15 fruits). **D)** Epicuticular wax composition and total wax load of ripe fruits (n=15 fruits). Data are presented as mean ± SEM. In A, the black dots represent individual data points. **E)** Simplified representation of cutin and wax biosynthesis and transport. Red letters indicate cuticle biosynthetic genes. **F)** RT-qPCR analysis of cuticle-related genes in fruit peels at the immature green stage (15 d post-anthesis). Heatmap shows log₂ fold changes (+FR vs. control, *n*=8). Asterisks denote significant difference between treatments (Student’s t-test, \**p*<0.05, \*\**p*<0.01, \*\*\**p*<0.001). The relative transcript values and gene abbreviations are detailed in Supplementary Table S3. Blue, yellow and green boxes indicate genes related to cutin, wax and transport/export, respectively, as represented in E. Abbreviations: ABCG, ABC transporter G family member; CUS, cutin synthase; FAE, fatty acid elongase multienzyme complex; GPAT, Glycerol-3-phosphate acyltransferase; KCS, beta-ketoacyl-CoA synthase; HCD, beta-ketoacyl-CoA reductase (KCR); beta-hydroxyacylCoA dehydratase; ECR, enoyl-CoA reductase; CER, fatty acyl-CoA reductase/ECERIFERUM; DCR, hydroxycinnamoyl transferase; WSD, wax synthase/diacylglycerol acyltransferase; MAH, mid-chain alkane hydroxylase; CYP, cytochrome P450, LTP(G), lipid transfer protein; VLC, very long chain.

Light microscopy analysis of outer pericarp sections from ripe fruits revealed significantly thicker cuticle layer in fruits from plants grown in FR-enriched conditions, including increments in cuticle thickness over the epidermal cells, and higher anticlinal peg depth and invagination degree compared to the control condition (Fig. 1A, B). In agreement, GC-MS quantification of cuticular components in ripe fruits revealed a promotive effect of low R/FR on the abundance of most cutin monomers, including the three major monomers 1,18-dioic- and 9,10,18-triOH-octadecanoic acids, and 9(10),16-diOH-hexadecanoic acid, leading to an increase of approximately 40% in total cutin load compared to the control (Fig 1C, Supplementary Table S2). FR-enriched conditions also promoted the deposition of epicuticular waxes, including increments of several *n*-alkanes and all triterpenoids, resulting in total wax load values approximately 20% higher than those of control counterparts (Fig. 1D; Supplementary Table S2).

Cutin monomers and wax compounds that form the cuticle layer are synthesized from activated FAs (Lara et al., 2015). In tomato fruits, cutin deposition peaks at the immature green (IG) stage (between 10 and 25 dpa, days post anthesis) (Reynoud et al., 2022; Mintz-Oron et al., 2008). Based on an extensive literature review, key tomato genes involved in fruit cuticle formation were selected for RT-qPCR analysis in peel isolates of IG (15 dpa) fruits. Accession numbers and supporting references for each selected gene are provided in Supplementary Table S1. Data revealed that genes encoding cytochrome P450 (*SlCYP86A69* and *SlCYP77A*), which catalyze the oxidation of activated FAs in the cutin biosynthesis pathway, were up-regulated in response to low R/FR conditions (Fig. 1E, F; Supplementary Table S3). Similarly, FR-enriched conditions also induced the expression of *SlGPAT6,* which encodes for glycerol-3-phosphate acyltransferase involved in the esterification step that generates 10,16-diOH C16-glycerol (Petit et al., 2021), and *SlDCR* (*DEFECTIVE IN CUTICULAR RIDGES*), a member of the BAHD family of acyltransferases required for cutin oligomer/polymer formation (Panikashvili et al., 2009), as well as GDSL esterase/lipase-enconding genes (*SlGDSL*) (Fig. 1E, F).

Genes encoding enzymes involved in wax biosynthesis were similarly up-regulated in FR-enriched conditions, such as *SlCER6 (ECERIFERUM6),* which encodes a beta-ketoacyl-coenzyme involved in the fatty acid elongation (FAE) cycle, whereas *SlKCS10* (*3-KETOACYL-CoA SYNTHASE10*) remained unchanged (Fig. 1E, F). Genes involved in the downstream steps that ultimately produce alkanes, aldehydes, alcohols, and esters, such as *SlCER26, SlCER1.5*, and *SlWSD1 (WAX SYNTHASE1)*, were also induced under low R/FR conditions. Except for *SlLTPG1 (GLYCOSYLPHOSPHATIDYLINOSITOL-ANCHORED LIPID PROTEIN TRANSFER2)*, all the analyzed genes involved in cutin monomer and wax transport were induced under FR-enriched conditions, including *SlABCG11*, *SlABCG32*, *SlABCG36*, *SlLTP1* and *SlLTPG2* (Fig. 1F).

Together, these results indicate that cutin and cuticular wax production and deposition in tomato fruits are promoted under simulated proximity shade conditions by FR-enriched light.

### Phytochrome B-dependent light perception negatively regulates cuticle-related gene expression and reduces cuticle formation in tomato fruits

Since PHYB plays a central role in transducing variations in the R/FR ratio into plant responses (Casal et al., 2012), we next explored the involvement of both tomato *PHYB* paralogs (*SlPHYB1* and *SlPHYB2,* Carlson et al., 2020; Pratt et al., 1995) in tomato cuticle metabolism.

Light microscopic imaging performed in ripe fruits of transgenic tomato plants overexpressing the constitutively active *SlPHYB1^Y253H^* (*SlYHB1^OE^*) and *SlPHYB2^Y252H^* (*SlYHB2^OE^*) alleles revealed significantly thinner cuticles over the epidermal cell walls and less developed anticlinal pegs between epidermal cells compared to the WT counterparts (Fig. 2A, B). Moreover, the cuticle invagination (i.e., how cutinized are the anticlinal and periclinal inner walls of the epidermal cells), which is of significant size in tomato fruits (España et al., 2014a), was conspicuously reduced in both transgenic lines (Fig. 2A, B). In agreement, fruits from S*lYHB1^OE^* and *SlYHB2^OE^* displayed lower total cutin load due to a reduction in virtually all major and minor cutin monomers (Fig. 2C, Supplementary Table S4). Additionally, fruits from both transgenic lines showed significantly lower accumulation of *n-*alkanes, branched alkanes, triterpenoids and total wax load, including a 1.8-fold reduction in the predominant wax *n*-alkane (C31), and reductions in several other minor compounds of cuticle wax compared to the WT (Fig. 2D, Supplementary Table S4). The marked decrease in cutin and epiculticular wax load triggered by *SlYHB1* and *SlYHB2* overexpression was explained by a global down-regulation of cuticle biosynthetic genes in peel isolates, including key genes involved in cutin monomer biosynthesis (*SlCYP86A69* and *SlDCR*) and polymerization (*SlCUS1, SlGDSL*), wax biosynthesis (*SlCER6*, *SlCER3.2*, *SlWSD1*) and lipid transport across the cell membrane (*SlLTP1*, *ABCG*s) (Fig. 2E, Supplementary Table S5).

**Figure 2.**
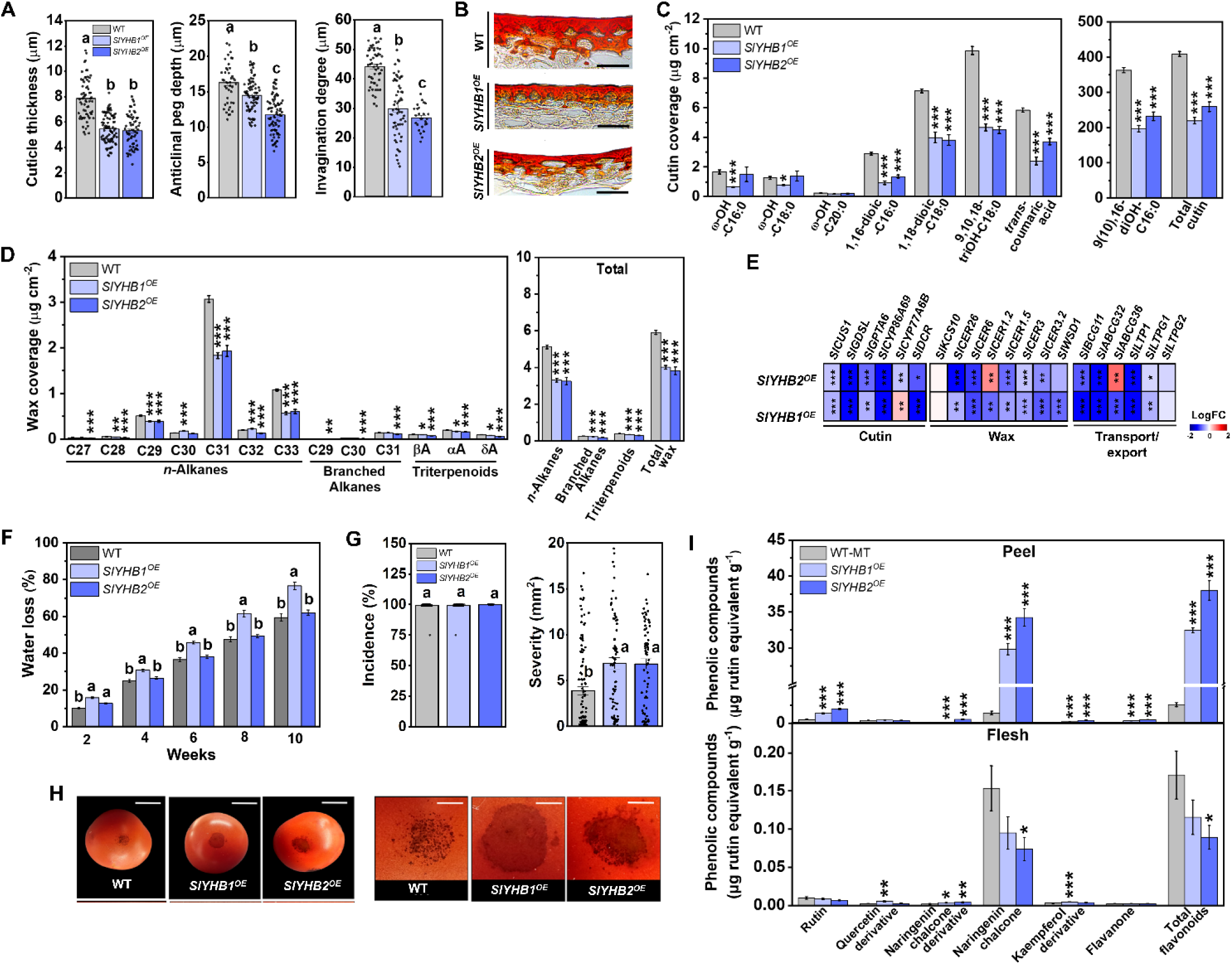
Overexpression of the constitutively active missense alleles *SlYHB1* and *SlYHB2* negatively regulates cuticle formation in tomato fruits, leading to higher susceptibility to fungal infection and enhanced postharvest water loss. Measurements were performed in fruits from wildtype (WT, cv. MicroTom) and *SlYHB1*- and *SlYHB2-* overexpressing (*SlYHB1^OE^* and *SlYHB2^OE^*, respectively) plants **A)** Cuticle thickness, anticlinal peg depth and invagination degree of ripe fruits (7 d after Breaker stage) (n=5 fruits). **B)** Representative light micrographs of pericarp cross-sections stained with Sudan-IV. Scale bars: 50 μm. **C)** Cutin monomer composition, total cutin load of ripe fruits (*n*=15 fruits). **D)** Epicuticular wax composition and total wax load of ripe fruits (n=15 fruits). **E)** RT-qPCR analysis of cuticle-related genes in fruit peels at Immature green stage (15 d post-anthesis). Heatmap shows log₂ fold changes (transgenic lines *vs*. WT, *n*=8). The relative transcript values and gene abbreviations are detailed in Supplementary Table S5. **F)** Percentage of water loss in ripe fruits across 10 weeks of storage. **G)** Disease incidence (%) and severity (lesion area in mm²) in ripe fruits after 72 h of inoculation with *Botrytis cinerea*. **H)** Representative necrotic lesions caused by the pathogen. Scale bars: 10 mm (left panels), 2 mm (right panels). **I)** Flavonoid profiles in peel (top) and flesh (bottom) tissues of ripe fruits (*n*=8 fruits). Data are presented as mean ± SEM. In A, the black dots represent individual data points. In A, F and G, different letters indicate statistically significant differences among genotypes (Tukey’s test, *P*<0.05). In C-E and I, asterisks denote significant difference in comparison to the WT (Student’s t-test, \**p*<0.05, \*\**p*<0.01, \*\*\**p*<0.001).

Modifications in the structure and/or composition of the cuticle can significantly impact postharvest aspects, altering water loss rates and susceptibility to pathogen infection (Arya et al., 2021). Despite the marked reductions in *n-*alkanes and cutin load in both transgenics, postharvest water loss was only altered in *SlYHB1^OE^* fruits, which exhibited approximately 25% more postharvest water loss rates than the WT counterparts after ten weeks of storage (Fig. 2F). Due to their drastically reduced cuticle deposition, ripe fruits of *SlYHB1^OE^* and *SlYHB2^OE^* were challenged with *B. cinerea*. To evaluate the role of the cuticle in this defense response, the spores were inoculated directly onto the sterilized, non-punctured surface of the fruits. After 72 h post-inoculation, the disease incidence (*i.e.,* percentage of affected lesions) was close to 100% for all genotypes, whereas the severity (*i.e.,* lesion area) was approximately 2-fold higher in *SlYHB1^OE^* and *SlYHB2^OE^* fruits than in the WT counterparts (Fig. 2G, H).

Besides being important nutraceutical compounds with antioxidant activity, flavonoids also play a significant role in postharvest disease resistance (Treutter, 2006). Flavonoid profiling in peel samples revealed increments of up to 40-fold in naringenin chalcone and total flavonoids in *SlYHB1^OE^* and *SlYHB2^OE^* ripe fruits compared to the WT. In contrast, *SlYHB1* and *SlYHB2* overexpression failed to promote flavonoid accumulation in the flesh of the ripe fruits (Fig. 2I), indicating the occurrence of differential regulatory mechanisms controlling flavonoid biosynthesis in inner and epidermal tomato fruit tissues.

The role of SlPHYB on fruit cuticle metabolism was also investigated by comparing single and double *Slphyb1* and *Slphyb2* mutants. Compared to the WT, limited anatomical differences in the fruit cuticle of single mutants were observed, except for a slight increase in the cuticle thickness in *Slphyb1* and approximately 10% reduction in the cuticle invagination degree in both single mutants. Interestingly, almost a 2-fold increase in the cuticle thickness over epidermal cells and highly cuticularized anticlinal pegs cuticle were detected in the *Slphyb1b2* double mutant (Fig. 3A, B).

**Figure 3.**
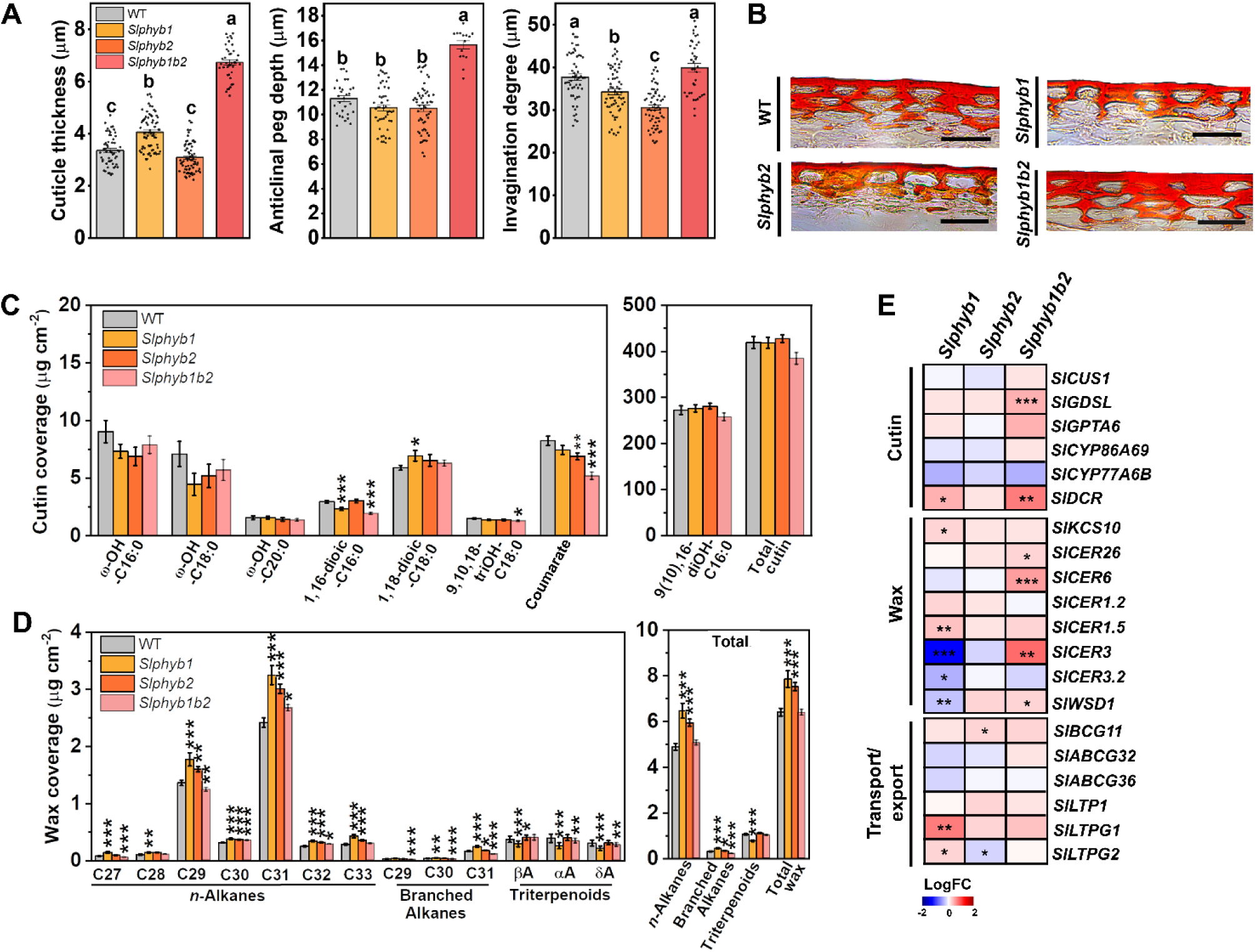
Loss of *SlphyB* function induces cuticle biosynthesis and deposition in tomato fruits. Measurements were performed in fruits from the wildtype (WT, cv. Moneymaker) and single and double mutants *Slphyb1, Slphyb2,* and *Slphyb1b2*. **A)** Cuticle thickness, anticlinal peg depth and invagination degree of ripe fruits (7 d after Breaker stage) (*n*=5 fruits). **B)** Representative light micrographs of pericarp cross-sections stained with Sudan-IV. Scale bars: 50 μm. **C)** Cutin monomer composition, total cutin load of ripe fruits (*n*=15 fruits). **D)** Epicuticular wax composition and total wax load of ripe fruits (*n*=15 fruits). **E)** RT-qPCR analysis of cuticle-related genes in fruit peels at the immature green stage (15 d post-anthesis). Heatmap shows log₂ fold changes (transgenic lines *vs*. WT, *n*=8). The relative transcript values and gene abbreviations are detailed in Supplementary Table S7. Data are presented as mean ± SEM. In A, the black dots represent individual data points, and different letters indicate statistically significant differences among genotypes (Tukey’s test, *P*<0.05). In C-E, asterisks denote significant difference in comparison to the WT (Student’s t-test, \**p*<0.05, \*\**p*<0.01, \*\*\**p*<0.001).

Chemical profiling of cuticle components revealed no significant changes in the abundance of 9(10),16-diOH-hexadecanoic acid and total cutin load across the genotypes (Fig. 3C, Supplementary Table S6), which might be explained by the limited changes in the cuticle invagination degree in the single and double mutants (Fig. 3A). In contrast, a clear induction of fruit cuticular wax deposition was observed in both single mutants, with increments between 20-30 % in the levels of total *n-*alkanes and total wax load compared to the WT counterparts (Fig. 3D, Supplementary Table S6). Single and double mutants showed increased expression of *SlKCS10*, and individually exhibited higher expression of nearly all analyzed genes, except for *SlLTP1, SlABCG32* and *SlABCG36* (Fig. 3E, Supplementary Table S7).

Collectively, these findings suggest some level of functional redundancy for both tomato *PHYB* paralogs in connecting light perception and fruit cuticle deposition by controlling the transcript abundance of key genes involved in cutin monomer biosynthesis, cutin polymerization and cuticular wax biosynthesis and transport.

### SlPIF3, SlPIF4 and SlBBX28 play a key role in light-regulated fruit cuticle formation

Photomorphogenesis-related transcription factors, such as PIFs and BBXs, have been increasingly implicated in controlling tomato fruit development and metabolism (Lira et al., 2020; Rosado et al., 2016; Shiose et al., 2024; Zhang and Lin, 2017). Among them, SlPIF3 and SlPIF4 regulate metabolic pathways impacting the fruit nutraceutical composition (Gramegna et al., 2019; Rosado et al., 2019), and SlBBX28 regulates flowering and fruit production (Lira et al., 2022). Analysis performed in leaves and ripening fruits of *SlYHB1^OE^* and *SlYHB2^OE^* plants revealed a drastic downregulation of *SlPIF3*, *SlPIF4* and *SlBBX28*, reinforcing the role for these TFs in SlPHYB signaling cascades (Fig. 4A).

**Figure 4.**
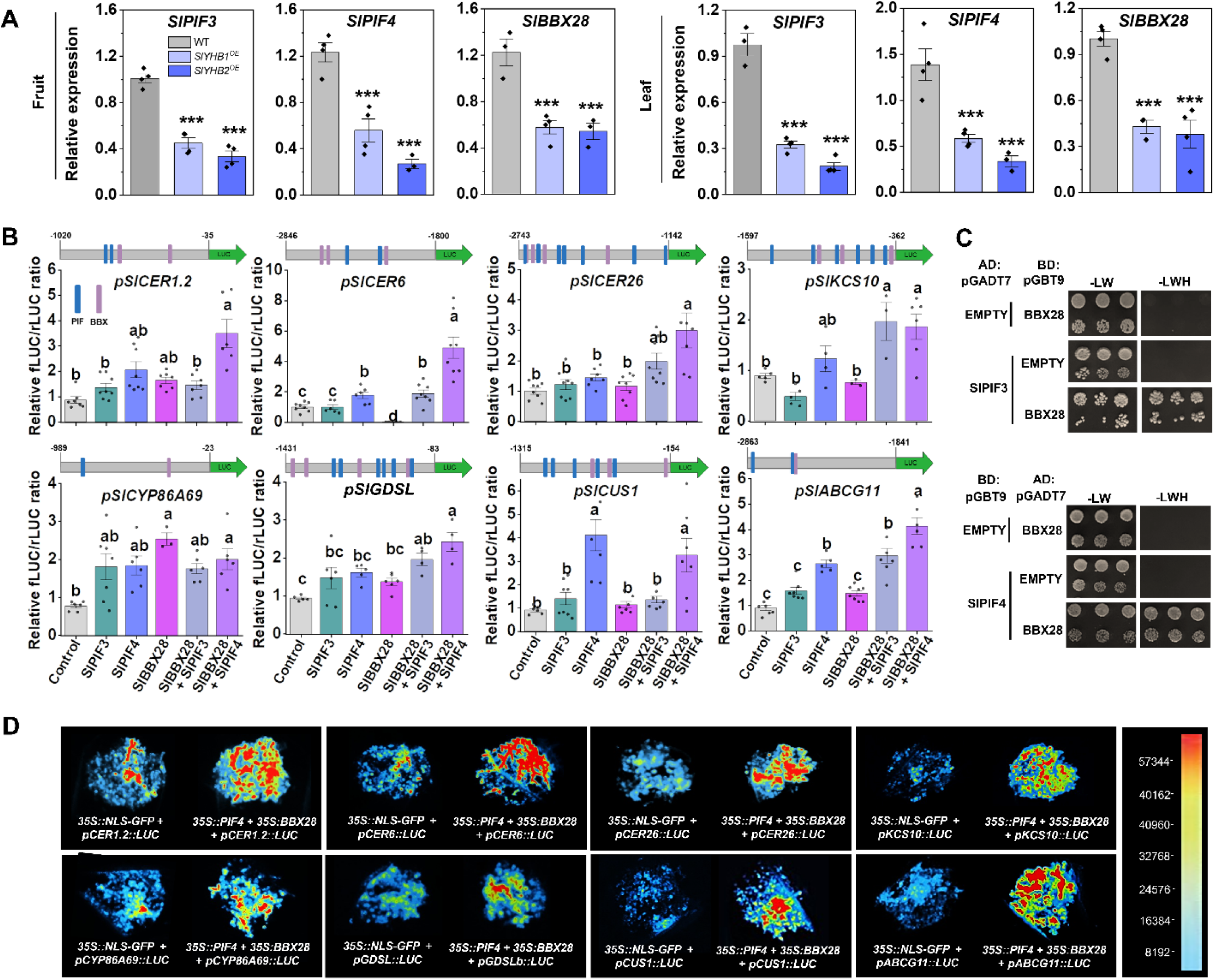
SlPIF3, SlPIF4 and SlBBX28 synergically activate promoters of key cuticle-related genes. **A)** RT-qPCR-based analysis of the transcript abundance of *SlPIF3, SlPIF4* and *SlBBX28* in immature green fruits and source leaves of wildtype (WT, cv. MicroTom) and *SlYHB1^OE^* and *SlYHB2^OE^* plants (*n*=8). Values are means ± SEM, and asterisks denote significant difference in comparison to the WT (Student’s t-test, \**p*<0.05, \*\**p*<0.01, \*\*\**p*<0.001). **B)** Transactivation expression assay in *Nicotiana tabacum* BY-2 protoplast cells showing activation of the promoters of key cuticle-related genes by SlPIF3, SlPIF4, SlBBX28, or their combinations. Luciferase activity is expressed as the ratio of firefly (fLUC) to Renilla (rLUC) luciferase relative to the negative control. Above each graph, schematic representations of the promoter regions fused to luciferase are shown (gray rectangle: promoter; green arrow: luciferase). Numbers indicate nucleotide positions upstream of the ATG. Blue and purple lines indicate PIF and BBX transcription factor binding motifs identified in each promoter, BBX: CCAAT (Ben-Naim et al., 2006), CCACA (Gnesutta et al., 2017); and PIF: E-box (CANNTG, Zhang et al., 2013). Data are presented as mean ± SEM with individual data points. Different letters indicate statistically significant differences among genotypes (Tukey’s test, *P*<0.05). **C)** Yeast two-hybrid interactions between SlBBX28 and SlPIF3 (top) or SlPIF4 (bottom). SlBBX28 was fused to the activation domain (AD) and SlPIF3 and SlPIF4 to the binding domain (BD). EMPTY, autoactivation control; -LW, positive control in non-selective medium without leucine and tryptophan; -LWH, selective medium without leucine, tryptophan, and histidine. Black boxes show three individual colonies at 10- and 100-fold dilutions. **D)** *Nicotiana benthamiana* leaves co-infiltrated with *Agrobacterium* containing luciferase reporter constructs driven by the promoter of cuticle-related genes, and *p35S::NLS-GFP* (control). Pseudocolors show the luminescence intensity (left).

*In silico* analysis revealed putative PIF- and BBX-binding motifs in the promoters of key genes involved in cutin monomer biosynthesis (*SlGDSL, SlCYP86A69*) and polymerization (*SlCUS1*), cuticular wax production (*SlCER1.2, SlCER6*, *SlCER26*, *SlKCS10*), as well as cutin monomer and wax transport across the membrane (*SlABCG11*) (Fig. 4B). Therefore, we next performed a dual-luciferase TEA in which reporter constructs of *fLUC* driven by the promoter regions of these genes were co-expressed in BY-2 protoplasts alongside effector constructs containing the coding sequences of *SlPIF3, SlPIF4 or SlBBX28*. Whereas SlPIF3 alone failed to transactivate the tested promoters, data showed that SlPIF4 alone can transactivate the *pSlCER6*, *pSlABCG11,* and particularly the *pSlCUS1* promoters (Fig. 4B). SlBBX28 alone exhibited a dual role, transactivating *pSlCYP86A69* while repressing *pSlCER6*, suggesting promoter-specific transcriptional modulation. As Y2H assays revealed that SlBBX28 physically interacts with both SlPIF3 and SlPIF4 (Fig. 4C), the combination of SlPIF3-SlBBX28 and SlPIF4-SlBBX28 as effectors was also tested in TEA assays (Fig. 4B). Compared to SlPIF3 alone, the SlPIF3-SlBBX28 combination activated a larger number of promoters: *pSlCER6, pSlGDSL, pSlKCS10* and *pSlABCG11* (Fig. 4B). Remarkably, promoters of all cuticle-related genes tested were transactivated by the SlPIF4-SlBBX28 combination, with an induction of up to 3-fold in their activities. Co-infiltration assays performed in *Nicotiana benthamiana* leaves further confirmed the intense induction of all promoters by the SlPIF4-SlBBX28, as evidenced by strong luminescence signals captured in the presence of both effectors (Fig. 4D).

Next, the roles of these TFs in tomato fruit cuticle formation were functionally assessed by using RNAi-mediated knockdown tomato lines *SlPIF3*^RNAi^, *SlPIF4^RNAi^* and *SlBBX28*^RNAi^. Light microscopic imaging of outer pericarp sections from ripe fruits revealed that both *SlPIF4* and *SlPIF3* silencing resulted in thinner cuticle layers over the epidermal cells and reduced the cuticle invagination degree of cuticle material (Fig. 5A, B). In agreement, the abundance of several cutin monomers and the total cutin load were significantly reduced in fruits of the *SlPIF3-* and *SlPIF4*-silenced lines (Fig. 5C, Supplementary Table S8). On the other hand, fruits from the *SlBBX28^RNAi^* line displayed slightly more cuticularized anticlinal pegs and reduced cuticle invagination degree compared to the WT (Fig. 5A).

**Figure 5.**
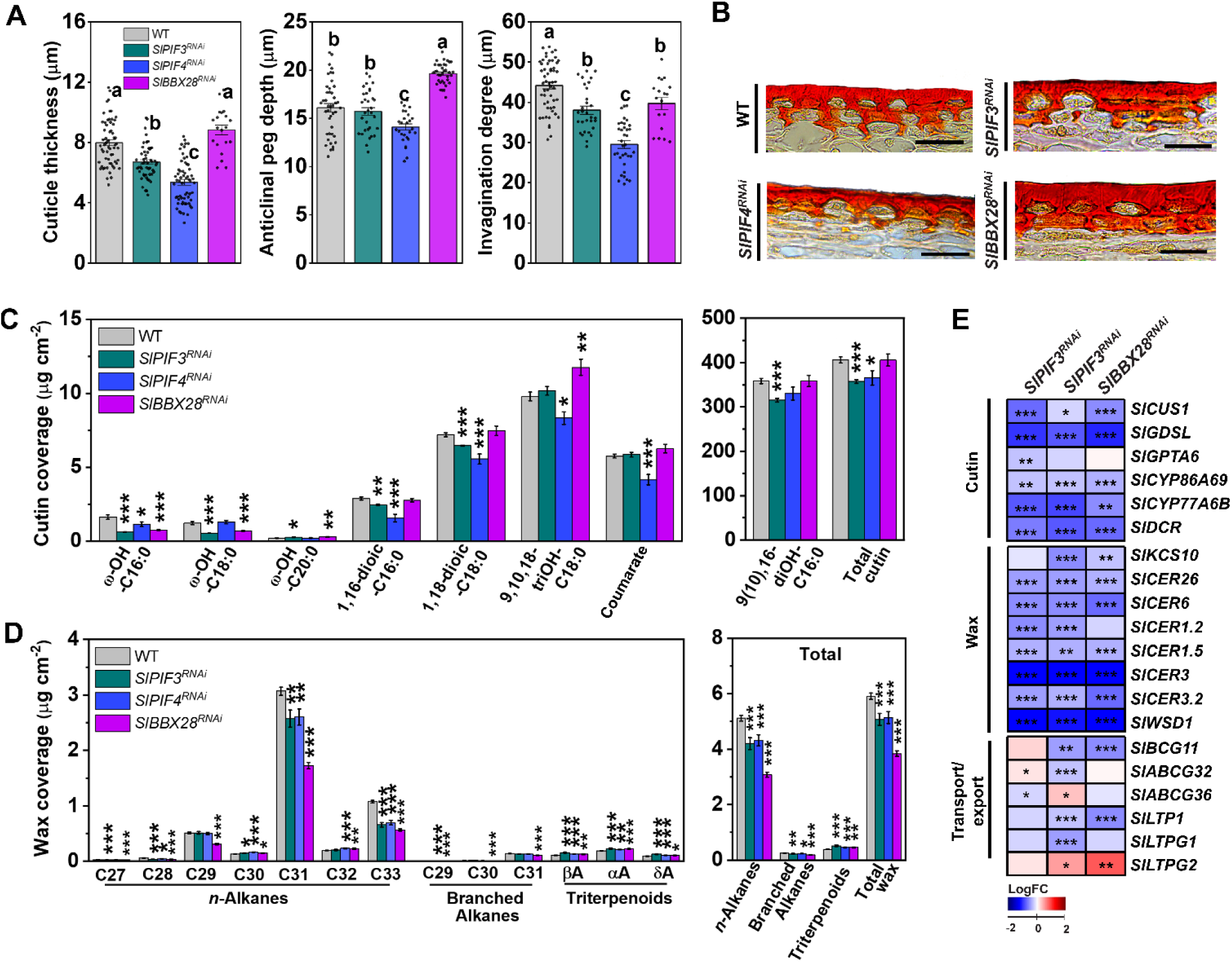
Silencing of *SlPIF3, SlPIF4* and *SlBBX28* represses cuticle biosynthesis and deposition in tomato fruits. Measurements were performed in fruits from wildtype (WT, cv. MicroTom) and RNA interference (RNAi)-mediated knockdown of *SlPIF3*, *SlPIF4* and *SlBBX28* (*SlPIF3^RNAi^, SlPIF4^RNAi^* and *SlBBX28 ^RNAi^*, respectively). **A)** Cuticle thickness, anticlinal peg depth and invagination degree of ripe fruits (7 d after Breaker stage) (*n*=5 fruits). **B)** Representative light micrographs of pericarp cross-sections stained with Sudan-IV. Scale bars: 50 μm. **C)** Cutin monomer composition, total cutin load of ripe fruits (*n*=15 fruits). **D)** Epicuticular wax composition and total wax load of ripe fruits (*n*=15 fruits). **E)** RT-qPCR analysis of cuticle-related genes in fruit peels at immature green stage (15 d post-anthesis). Heatmap shows log₂ fold changes (transgenic lines *vs.* WT, *n*=8). The relative transcript values and gene abbreviations are detailed in Supplementary Table S5. Data are presented as mean ± SEM. In A, the black dots represent individual data points, and different letters indicate statistically significant differences among genotypes (Tukey’s test, *P*<0.05). In C-E, asterisks denote significant difference in comparison to the WT (Student’s t-test, \**p*<0.05, \*\**p*<0.01, \*\*\**p*<0.001).

A consistent negative impact on epicuticular wax deposition was observed in response to *SlPIF3, SlPIF4*, and *SlBBX28* silencing, with reductions in virtually all *n*-alkanes, branched alkanes, and triterpenoids (Fig. 5D, Supplementary Table S8). Compared to the WT, reductions of approximately 30% in total n-alkanes and wax load were observed for *SlBBX28^RNAi^* fruits, whereas *SlPIF3* and *SlPIF4* silencing resulted in a decline of about 20% in these parameters.

With very few exceptions, the transcript abundance of key cutin biosynthetic genes in the fruit peel tissues of all three silenced lines was drastically reduced, including ∼2-fold downregulation of *SlGDSL, SlCYP86A69, SlCYP77A6B* and *SlDCR* in the transgenics compared to the WT (Fig. 5E, Supplementary Table S5). All cuticular wax biosynthetic genes analyzed were drastically reduced in all three silenced lines (Fig. 5E).

Therefore, SlPIF3, SlPIF4, and SlBBX28 seem to act as positive regulators in the SlPHYB-dependent regulation of tomato fruit cuticle formation through the direct induction of genes involved in multiple key steps of cutin and wax biosynthesis.

### Transcriptional reprogramming behind the PHYB-dependent regulation of fruit cuticle biosynthesis

Transcriptomic analysis was performed to further elucidate the mechanisms behind the PHYB-PIF-BBX-dependent regulation of tomato fruit cuticle biosynthesis. The RNA-seq profiling included peel isolates of IG fruits from WT plants cultivated under control and FR-enriched conditions, as well as single and double *Slphyb* mutants and *SlYHB1^OE^*, *SlYHB2^OE^*, *SlPIF3^RNAi^*, *SlPIF4^RNAi^* and *SlBBX28 ^RNAi^* transgenic lines under normal light conditions (Fig. 6).

**Figure 6.**
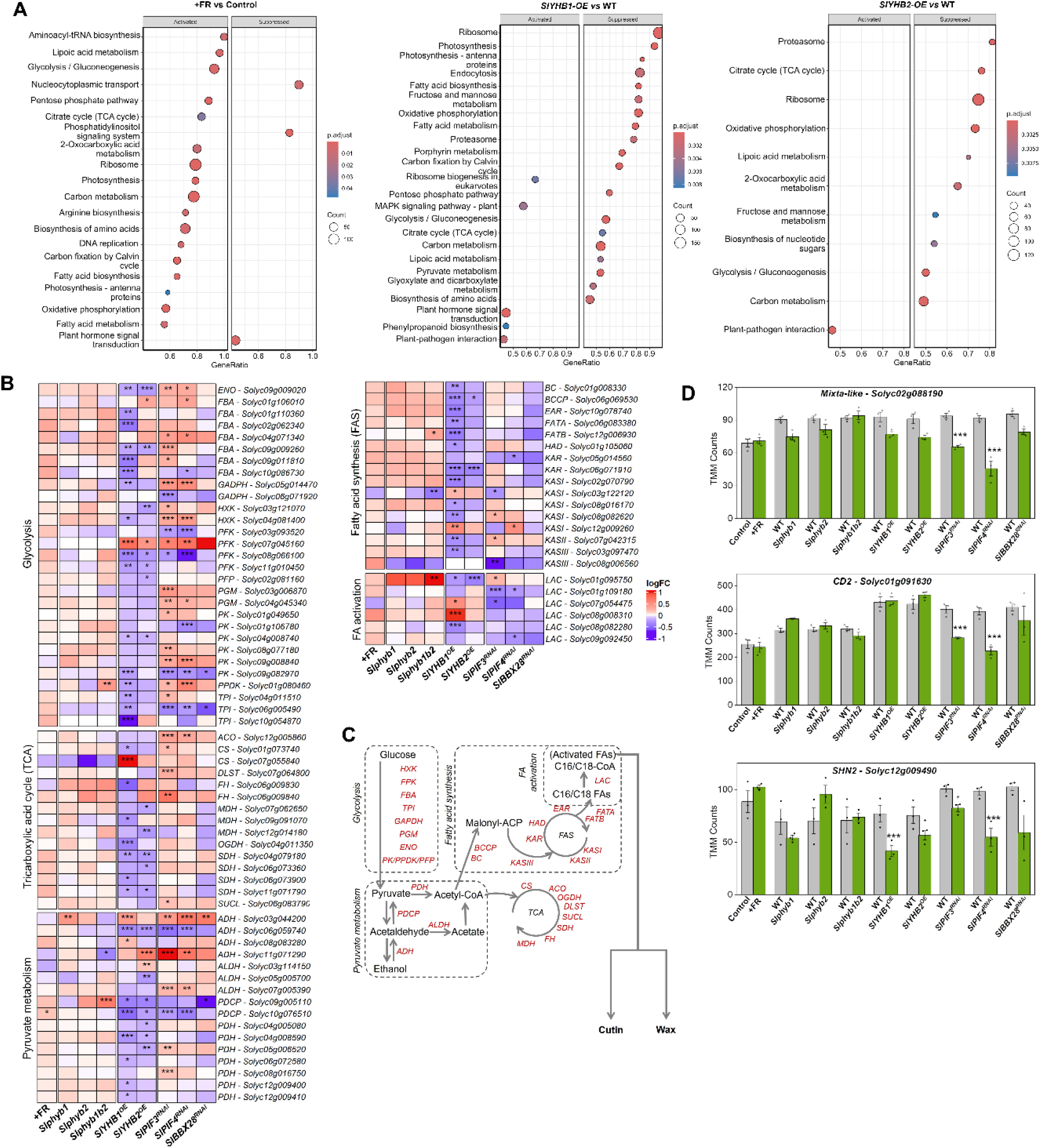
Phytochrome B-mediated signaling induces transcriptional reprogramming of core metabolism in tomato fruit peel. RNA-seq profiling of peel isolates of fruits at the immature green stage (15 d post-anthesis, *n*=3) from wildtype (WT) plants cultivated under control and far-red-enriched conditions (+FR), as well as single and double *Slphyb* mutants (*Slphyb1, Slphyb2*, and *Slphyb1b2*) and *SlYHB1^OE^, SlYHB2^OE^, SlPIF3^RNAi^, SlPIF4^RNAi^ and SlBBX28^RNAi^* transgenic lines under normal light conditions. **A)** Dot plot of significantly enriched KEGG pathways that are either activated or suppressed in +FR *vs*. control, *SlYHB1^OE^ vs*. WT, and *SlYHB2^OE^ vs*. WT (left to right). Dot size represents the number of genes in each pathway, and gradient color indicates adjusted p-value. **B)** Expression profile of differentially expressed genes involved in core metabolism pathways. Heatmap shows log₂ fold changes (+FR *vs*. control or transgenic/mutant lines *vs*. WT). Transcript values and abbreviations are described in Supplementary Table S10. **C)** Schematic representation of modulated genes (highlighted in red) from metabolic pathways shown in B. **D)** TMM-normalized expression values of transcriptional regulators involved in cuticle biosynthesis in tomato: *SlCD2* (*CUTIN DEFICIENT2*)*, SlMIXTA-Like* and *SlSHN2* (*SHINE2*). Data are presented as mean ± SEM. In B and C, asterisks denote significant difference in comparison to the WT or control condition (Student’s t-test, \**p*<0.05, \*\**p*<0.01, \*\*\**p*<0.001).

Enrichment analysis of KEGG identified metabolic pathways critical for cutin and wax biosynthesis, such as FA biosynthesis and metabolism, as induced in fruits from plants grown under low R/FR (Fig. 6A). Upstream pathways related to primary carbon fixation and central carbon processing were also activated under simulated shade conditions, including the carbon metabolism, glycolysis, pyruvate metabolism and the tricarboxylic acid cycle (TCA) pathways. In contrast, *SlYHB1*^OE^ and *SlYHB2*^OE^ exhibited widespread suppression of these same pathways (Fig. 6A). In agreement, fruits from the *Slphyb* single and double mutants showed activation of lipid-related pathways similar to those observed in the low R/FR plants (Supplementary Table S9).

Based on these findings, we further examined the expression profiles of DEGs within KEGG-enriched pathways associated with the metabolism of activated FAs, the precursors for cuticle formation. DEGs related to upstream routes such as glycolysis, pyruvate metabolism, the TCA cycle, and FA synthesis and activation were massively downregulated in *SlYHB1^OE^* and *SlYHB2^OE^*, with a tendency of repression also in *SlPIF3^RNAi^*, *SlPIF4^RNAi^* and *SlBBX28^RNAi^* fruits (Fig. 6B). Therefore, an overall downregulation of different metabolic pathways required for the production of precursors for cutin and wax biosynthesis was observed in all transgenic lines (Figure 6B, C). In contrast, these same genes were only marginally affected in response to simulated shade or in the *Slphy* single and double mutants (Fig. 6B, Supplementary Table S10).

Besides these changes in key metabolic routes, an additional mechanism behind SlPHYB-dependent regulation of cuticle metabolism seems to involve the modulation of TFs previously identified as regulators of cuticle formation in tomato fruits (reviewed by Petit et al., 2021). For example, *SlMIXTA-like* and *CUTIN DEFICIENT2* (*SlCD2*), which encodes positive transcriptional regulators of cuticle biosynthesis (Isaacson et al., 2009; Lashbrooke et al., 2015; Nadakuduti et al., 2012), were downregulated in *SlPIF3^RNAi^* and *SlPIF4^RNAi^* fruits (Fig. 6D). In addition, *SHINE2* (*SlSHN2*), which promotes cutin deposition and regulates epidermal patterning (Bres et al., 2022), was downregulated in *SlYHB1^OE^*, *SlYHB2^OE^*, *SlPIF3^RNAi^* and *SlPIF4^RNAi^* fruits (Fig. 6D).

Altogether, our findings indicate that changes in central carbon metabolism (*e.g.,* glycolysis and TCA) and plastidial *de novo* C16/C18 FA biosynthesis and FA activation, alongside the modulation of cutin-related TFs (*e.g., SlCD2*, *SlMIXTA-like* and *SlSHN2*), may also be implicated in PHYB-dependent regulation of cuticle biosynthesis in tomato fruits.

## DISCUSSION

Photoreceptors and downstream regulatory proteins have been increasingly identified as important regulators of metabolic pathways leading to the accumulation of specialized metabolites, particularly antioxidants, in fleshy fruits (Li et al., 2018). Our findings uncover a new facet of light signaling in fruit development, revealing that PHYB and downstream photomorphogenesis-related TFs form a molecular circuit regulating the biosynthesis and deposition of the cuticle, a crucial structure for fruit defense and extended shelf life.

Although still poorly characterized, evidence suggests PHY-mediated light perception as a negative regulator of cuticle formation in leaves and etiolated seedlings (Qiao et al., 2020; Ma et al., 2024). More specifically, etiolated *A. thaliana* seedlings expressing the constitutively active alleles *PHYB^Y276H^* and *PHYA^Y242H^* showed reduced expression of the cutin biosynthetic genes *LONG-CHAIN ACYL-CoA SYNTHETASE* 2 (*AtLACS2*)*, BODYGUARD 1* (AtBDG1) and

*AtDCR* under continuous dark; although the impact on epicuticular wax and cutin accumulation was not assessed (Ma et al., 2024). In addition, *Zmphyb1 Zmphyb2* double mutants displayed increased total alkanes and free fatty acids in maize leaves (Qiao et al., 2020).

In line with these reports, here, we reveal a key repressor role for both tomato *PHYB* paralogs in regulating both fruit cuticle composition and structure, as supported by converging evidence from *phyb* loss-of-function and constitutively active *PHYB* alleles. The *Slphyb1b2* double mutant showed increased cuticle thickness and anticlinal peg depth compared to single mutants, highlighting partial redundancy between the two paralogs in controlling fruit cuticle development (Fig. 3). Conversely, an overall downregulation of cutin and wax biosynthetic genes and reduction in cutin and epicuticular wax load were evidenced in *SlYHB1^OE^* and *SlYHB2^OE^* lines (Fig. 2).

PHYB is inactivated under low R/FR light ratios, a condition that prevails in shade environments and often results from surrounding vegetation (Casal, 2012; Legris et al., 2019). Simulated proximity shade, via FR-light enrichment without altering total PAR intensity, promoted effects consistent with the repressive role of SlPHYB1/B2, as fruits from plants grown under this condition exhibited increased cuticle thickness and higher accumulation of cutin monomers and wax compounds, accompanied by the upregulation of several cuticle-related genes (Fig. 1). This expands previous insights from vegetative tissues exposed to continuous light or dark treatments (Go et al., 2014; Hooker et al., 2002; Ma et al., 2024), revealing that R/FR fluctuations, detected by a PHYB-mediated pathway, confer an additional regulatory layer over cuticle deposition in fruits developing in an ever-changing light environment. Moreover, our findings also have important implications when comparing cuticle-related traits of fruits growing in the interior or exterior canopy, as significant differences in wax accumulation have been previously observed in fleshy fruits under these conditions (reviewed by Liu et al., 2023).

Strikingly, the SlPHYB1/B2-mediated repression of fruit cuticle deposition compromised its protective role during postharvest storage. A first outcome from this detrimental effect was the increased water loss in *SlYHB1^OE^* fruits (Fig. 2). Although both transgenic lines displayed comparable reductions in total epicuticular wax, the cutin layer, which comprises more than 90 % of the tomato fruit cuticle (Liu et al., 2023), was reduced to a greater extent in *SlYHB1^OE^* compared to *SlYHB2^OE^* fruits, likely explaining the different postharvest water loss rates observed in these transgenics. A second detrimental effect was the higher susceptibility of *SlYHB1^OE^* and *SlYHB2^OE^* fruits to *B. cinerea*, reflecting the key role played by the cuticle as a frontline barrier against pathogen invasion (Ziv et al., 2018). As our experimental setup involved the spore inoculation directly onto the intact fruit surface, we can conclude that such susceptibility to fungal infection was linked to the abnormal cuticle formation in these transgenic lines.

Relevant information about the light-dependent regulation of flavonoids throughout the fruit tissues was also obtained (Fig. 2). Adding to the previous observation that the fruit-specific overexpression of *SlYHB2* resulted in higher flavonoid levels in whole pericarp samples, including a mixture of both flesh and peel tissues (Alves et al., 2020), our tissue-specific approach reveals that this increase occurs exclusively in the peel of both *SlYHB1^OE^* and *SlYHB2^OE^* fruits. The presence of chlorophylls within the chloroplasts of green tomato fruits increasingly reduces the R/FR ratios as light penetrates through the pericarp (regarded as ‘self-shading’), which is proposed to limit the PHY-dependent induction of carotenoid biosynthetic genes in inner fruit tissues (Llorente et al., 2016). According to our findings, even the constitutive overexpression of the hyperactive *SlYHB1* and *SlYHB2* alleles completely failed to activate flavonoid biosynthesis in the inner fruit tissues, revealing that the reduced levels of these compounds in inner pericarp tissues may not be due to a self-shading effect. From the perspective of postharvest fruit deterioration, despite the well-known role of flavonoids in defense against necrotrophic pathogens in fruits (Ramaroson et al., 2022), even the 40-fold increment of these antioxidants in the peel of the transgenic lines was insufficient to compensate for the impaired cuticle, reinforcing the key role played by this first line of defense. Moreover, flavonoids have been increasingly shown to modify cuticle structure by interacting with its components, thus altering important cuticle properties (España et al., 2014a; Reynoud et al., 2021). Therefore, the significant induction of flavonoid biosynthesis in tomato fruit peel tissues triggered by PHYB signaling may also have contributed to changes in the cuticle properties observed in *SlYHB1^OE^* and *SlYHB2^OE^* fruits.

By exploring the molecular mechanisms underlying the PHY-mediated control of tomato fruit cuticle regulation, we identified SlPIF3, SlPIF4, and SlBBX28 as transcriptionally repressed in *SlYHB1*- and *SlYHB2*-overexpressing plants. These findings align with reports indicating that PIFs activate their own transcription (Shiose et al., 2024), and active PHYB can regulate PIF activity by promoting their proteasomal degradation and/or repressing their transcriptional activity (Chen and Chory, 2011; Park et al., 2012). In addition, the involvement of SlPIF3, SlPIF4, and SlBBX28 as positive regulators of tomato fruit cuticle development is supported by the fact that: (i) fruits from RNAi lines for these TF-encoding genes displayed reduced cuticle thickness and lower cutin and epicuticular wax load due to the downregulation of multiple cuticle-related genes, and (ii) both the isolated and combined action of SlPIF4 and SlBBX28, and to a lesser extent SlPIF3, transactivated promoters of key cutin and wax biosynthetic genes.

B-box domains enable transcriptional regulation either through direct promoter binding (Xu et al., 2016) or by modulating the transcriptional activity of other TFs via heterodimerization (Song et al., 2020; Tripathi et al., 2017). Indeed, we observed the physical interaction between SlBBX28 and SlPIF3/4, and while SlPIF4 and SlBBX28 alone were able to activate a subset of cuticle-related gene promoters, their effect was less pronounced compared to the stronger activation observed upon co-expression. Therefore, the regulation of wax and cutin biosynthetic gene expression in response to phyB-mediated detection of R/FR conditions in tomato fruits may occur either directly by SlPIF4 or SlBBX28, or more effectively through heterodimer formation between SlBBX28 and SlPIF3/4.

Our data also indicated that, besides regulating downstream TFs, SlPHYB1/B2 may influence fruit cuticle formation via changes in the central carbon metabolism of the developing fruits, including glycolysis and pyruvate metabolism, which supply acetyl-CoA for FA synthesis, as well as the TCA cycle, which contributes both to FA oxidation and the generation of biosynthetic precursors (Niehaus, 2021). This regulation extended through the plastidial *de novo* C16/C18 FA biosynthesis and FA activation (Fig. 6B, C), which ultimately form the C16/C18-CoA that feeds into the wax and cutin biosynthetic pathway (Fich et al., 2016; Schnurr et al., 2004). Besides providing energy for photosynthesis, light also provides key information for metabolic fluxes (e.g., carbon partitioning, the TCA cycle, and glycolysis, secondary metabolism) across the plant through dedicated signaling pathways (Han et al., 2017). Moreover, changes in PHY stationary state due to supplementation with FR radiation can enhance fruit sink strength in tomato, promoting fruit growth by enhancing dry mass allocation to these organs (Ji et al., 2020; Vincenzi et al., 2024, 2025). In agreement, the overexpression of the *SlYHB1* allele negatively impacts starch and soluble sugar accumulation in tomato fruits (Alves et al., 2020). Therefore, these multiple lines of evidence support our conclusion that PHYB-dependent downregulation of key fruit metabolic fluxes may further limit the supply of precursors for the cuticle assembly in developing tomato fruits.

Taking our findings together, we propose a regulatory pathway (Fig. 7) where SlPHYB1/B2 signaling represses cuticle deposition in fruits by: (i) antagonizing the activity of SlPIF3, SlPIF4, and SlBBX28, which act as positive regulators of cuticle formation, and (ii) transcriptionally reprogramming the fruit central carbon metabolism, thereby constraining the supply of C16/C18 FAs and CoA thioesters required for cuticle biosynthesis.

**Figure 7.**
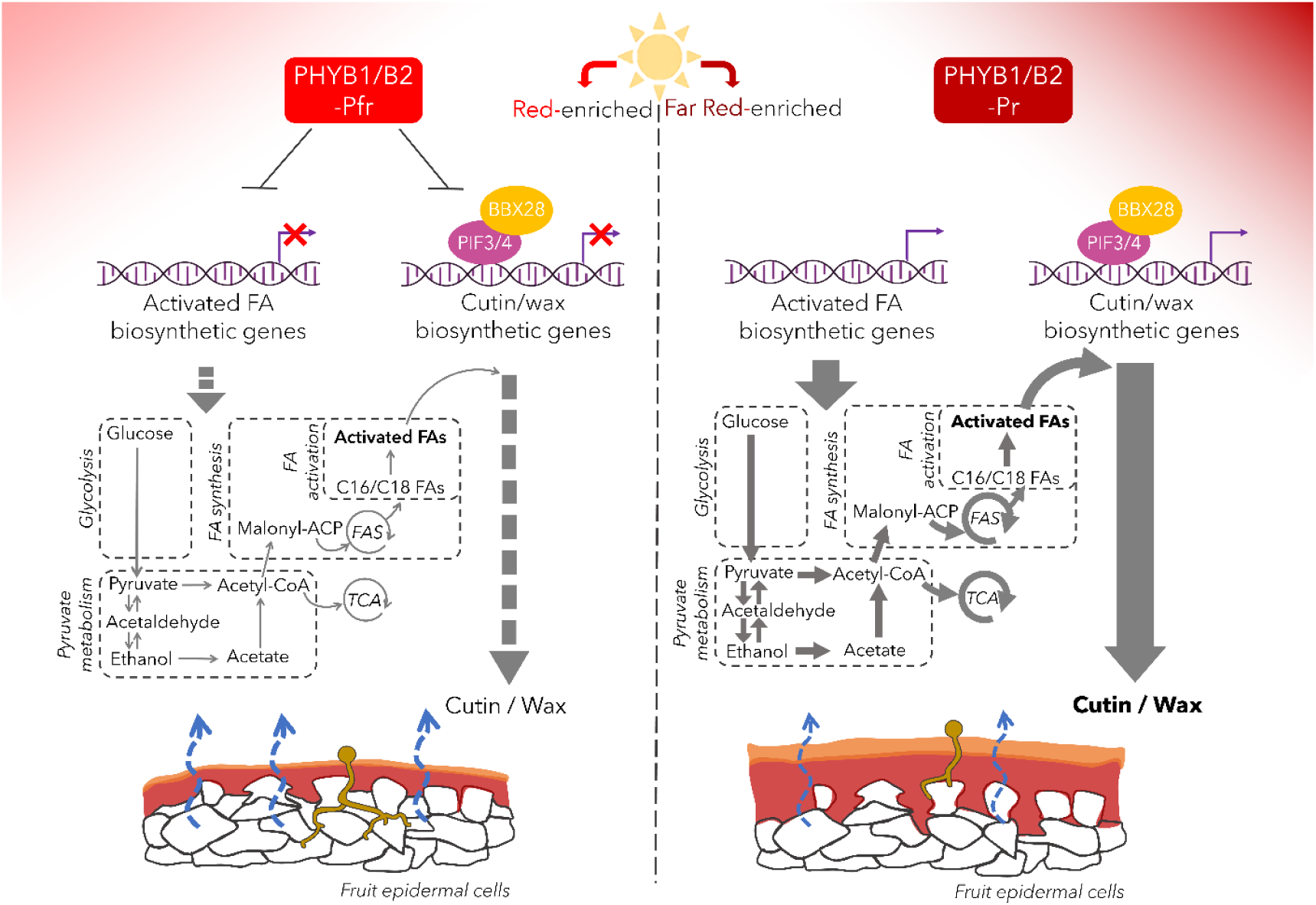
Proposed regulatory model for SlphyB1/B2-mediated control of fruit cuticle deposition. Upon red-light absorption, activated SlPHYB1/B2 (Pfr form) represses downstream positive regulators SlPIF3/4 and SlBBX28, which otherwise physically interact to activate the expression of cuticle-related genes. Concurrent downregulation of genes from pathways related to the biosynthesis and activation of fatty acids (FAs), likely reduces the pool of activated FAs feeding into cutin and wax biosynthetic pathways, thereby constraining cuticle formation and resulting in a thinner protective layer over fruit epidermal cells (left). By contrast, conversion of SlPHYB1/B2 to the inactive Pr form under far-red-enriched light conditions relieves this repression, leading to increased cuticle load (right). The red layer above epidermal cells represents polymerized cutin, the orange layer symbolizes epicuticular waxes, the yellow structure represents a germinated *Botrytis cinerea* spore, and the curved blue arrows represent loss of water across the cuticle. The thickness of gray arrows represents increased (thicker) or reduced (thinner) metabolic fluxes. Abbreviations: FAS, fatty acid synthase; TCA, tricarboxylic acid cycle.

Besides uncovering a novel regulatory circuit connecting light signaling and cuticle formation, our findings also highlight the importance of profiling a wide range of fruit quality traits in attempts to manipulate light signaling genes for boosting nutraceutical traits in fleshy fruits. The occurrence of similar trade-offs, where fruit biofortification may come at the cost of compromising fruit shelf life due to increased postharvest water loss and higher susceptibility to pathogens, remains an important topic for further investigation in tomatoes and other fleshy fruits.

## Acknowledgements

This work was supported by the São Paulo Research Foundation (FAPESP) (grants 23/08610-4; 23/03330-3; 21/10498-2, 21/13931-9, 24/13844-7, 24/02315-3), Conselho Nacional de Desenvolvimento Científico e Tecnológico (CNPq) (grant 304877/2022-0), and Coordenação de Aperfeiçoamento de Pessoal de Nível Superior – Brasil (CAPES) – Finance Code 001. We thank Dr. Deborah Yara Alves Cursino dos Santos (University of São Paulo) for helping during the GC-MS-based identification of the cutin monomers and wax components.

## Author contributions

LF and MR conceived the project and supervised the experiments; LDLGF, BO, JRM, PPAO and JNMU performed experiments; DD and CR provided technical assistance and contributed to the data analysis and discussion; SCSA and BO conducted bioinformatics analysis; LF, LDLGF, BO and MR wrote the article with contributions from other authors.

## Data availability

All data supporting the findings of this study are available within the paper and within its supplementary materials published online.

## Supplementary data

The following supplementary data are available.

**Supplementary Table S1**. Primer sequences used in this study and literature references for genes selected for RT-qPCR analysis.

**Supplementary Table S2**. Cutin and wax composition of fruit cuticles from control and shaded plants.

**Supplementary Table S3.** Transcript abundance of cuticle-related genes in fruit peels from control and shaded plants.

**Supplementary Table S4.** Cutin and wax composition of fruit cuticles from *SlYHB*-overexpressing plants.

**Supplementary Table S5.** Transcript abundance of cuticle-related genes in fruit peels from *SlYHB*-overexpressing and *SlPIF*/*SlBBX*-silenced plants.

**Supplementary Table S6.** Cutin and wax composition of fruit cuticles from *Slphy* mutants.

**Supplementary Table S7.** Cutin and wax composition of fruit cuticles from *Slphy* mutants.

**Supplementary Table S8.** Cutin and wax composition of fruit cuticles from *SlPIF*/*SlBBX*-silenced plants.

**Supplementary Table S9.** KEGG-based classification identified from transcriptome analysis.

**Supplementary Table S10.** Cuticle-related differentially expressed genes (DEGs) in fruit peel samples.

## Notes

### Competing Interest Statement

The authors have declared no competing interest.

